# Improving the performance of minimizers and winnowing schemes

**DOI:** 10.1101/104075

**Authors:** Guillaume Marçais, David Pellow, Daniel Bork, Yaron Orenstein, Ron Shamir, Carl Kingsford

## Abstract

The minimizers scheme is a method for selecting *k*-mers from sequences. It is used in many bioinformatics software tools to bin comparable sequences or to sample a sequence in a deterministic fashion at approximately regular intervals, in order to reduce memory consumption and processing time. Although very useful, the minimizers selection procedure has undesirable behaviors (e.g., too many *k*-mers are selected when processing certain sequences). Some of these problems were already known to the authors of the minimizers technique, and the natural lexicographic ordering of *k*-mers used by minimizers was recognized as their origin. Many software tools using minimizers employ ad hoc variations of the lexicographic order to alleviate those issues.

We provide an in-depth analysis of the effect of *k*-mer ordering on the performance of the minimizers technique. By using small universal hitting sets (a recently defined concept), we show how to significantly improve the performance of minimizers and avoid some of its worse behaviors. Based on these results, we encourage bioinformatics software developers to use an ordering based on a universal hitting set or, if not possible, a randomized ordering, rather than the lexicographic order. This analysis also settles negatively a conjecture (by Schleimer *et al*.) on the expected density of minimizers in a random sequence.

The software used for this analysis is available on GitHub: https://github.com/gmarcais/minimizers.git.

## 1 Introduction

The winnowing scheme was introduced by Schleimer *et al*. [2003] to fingerprint documents for plagiarism detection. Independently, the minimizers algorithm was introduced by Roberts *et al*. [2004b] to compute overlaps between sequencing reads. Even though these algorithms were designed for different purposes, they are essentially identical. The original minimizers scheme compares nucleotide *k*-mers using the lexicographic ordering, while the winnowing scheme hashes the *k*-grams of letters in a document, effectively randomizing the order.

In the bioinformatics field, minimizers have since been used in many different settings, such as binning input sequences [Roberts *et al*., 2004b,a, Li and Yan, 2015, Deorowicz *et al*., 2015], generating sparse data structures [Grabowski and Raniszewski, 2015, Ye *et al*., 2012], and sequence classification [Wood and Salzberg, 2014]. All these applications share the need for a small signature, or fingerprint, in order to recognize longer exact matches between sequences.

The winnowing scheme is defined as follows: given an input sequence *S* and parameters *k* and *w*, select the smallest *k*-mer (according to a predefined ordering) in each window of *w* consecutive *k*-mers in *S* (such a window contains *w* + *k* − 1 bases). In this paper, we will interchangeably use the terms “minimizers scheme” or “winnowing scheme”. Either term does not imply a particular ordering as we study the effect of various orderings on these schemes. The minimizers (or fingerprints), are, depending on the application, either the set of positions in *S* of the selected *k*-mers, or the set of selected *k*-mers itself. The terms fingerprint and minimizer are interchangeable in the remainder of this paper.

These schemes are designed to select a set of *k*-mers that is as sparse as possible while satisfying the following two properties. First, the sequence is approximately uniformly sampled; that is, the distance between two selected *k*-mers is always less than *w*. Second, if two sequences share an exact subsequence of length at least *k* + *w* – 1, then those two sequences have at least one minimizer in common.

Many of the tools mentioned above do not use lexicographic ordering of the *k*-mers, as originally proposed in Roberts et al. [2004b]. It was recognized early on that homo-polymer runs, particularly repeated As, can cause a lexicographic minimizer algorithm to select many consecutive *k*-mers as minimizers in genomics applications, leading to a high minimizer density. Because A is the smallest letter, if the 5-mer *AAAAC* is selected as a minimizer in a window, it is likely that the shifted 5-mer *AAAC*_*x*_ (for any base *x*) will also be selected as a minimizer in a subsequent window, and *AAC_xy_* in the following window, and so on.

Another problem with the lexicographic ordering is that any one of the 4^*k*^ possible *k*-mers can theoretically be chosen as a minimizer in some window. In particular, when *k* is short, many or even all possible *k*-mers could appear as a minimizer. If the minimizers are used for binning sequences, then potentially a very large number of bins will be necessary.

Schleimer et al. defined the *density* of a *k*-mer selection scheme for a given sequence *S* as the fraction of the *k*-mers that are selected. Formally, let *d(A, S, k)* denote the density of procedure *A* and *A(S, k)* as the set of selected *k*-mers positions by *A* on sequence *S*, then

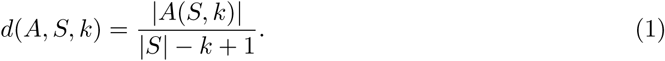

The *expected density* of procedure *A, d(A, k)*, is calculated by taking the expectation over all possible sequences *S*, with every base chosen independently with equal probability.

Schleimer et al. show that the winnowing scheme with a random ordering has expected density of 2/(*w* + 1). In practice, the minimizers scheme with lexicographic ordering has density greater than 2/(*w* + 1). In addition, they define a *local scheme* as a procedure that only has access to the sequence within a given window when selecting a *k*-mer. In other words, the *k*-mer selected from a window can be expressed as a function of only the identity and relative ordering of the bases within the window itself, and does not depend on the content of any other window or on the position of the window within the overall sequence. They prove that the expected density of a local scheme has a lower bound of 1.5/(*w* + 1), but they conjecture that this bound cannot be achieved in practice, and that the true lower bound is in fact 2/(*w* +1). This conjecture would therefore imply that the winnowing scheme with random order is an optimal local scheme.

In this paper, we investigate the effect of different orderings of the *k*-mers on the density of the winnowing scheme. By using universal *k-mer hitting sets*, as defined in Orenstein et al. [2016], we show how to create orderings that lead to densities smaller than with the lexicographic or randomized ordering. The small universal hitting sets created by the DOCKS algorithm developed in Orenstein et al. [2016] achieve densities below 1.8/(*w* + 1). As a consequence, the above conjecture of Schleimer et al. does not hold and densities below 2/(*w* + 1) are achievable with a local scheme. Using some properties of the universal *k*-mer hitting sets, we also show that, surprisingly, the winnowing scheme with random ordering can itself have a density slightly below 2/(*w* + 1). We also explain why the original minimizers procedure, using the lexicographic order, performs worse than random ordering.

Finally, we look at the potential effect of using universal hitting set orderings on bioinformatics applications. In the case of DNA sequence binning, such as performed by the *k*-mer counters KMC2 [Deorowicz *et al*., 2015] and MSPKmerCounter [Li and Yan, 2015], we compare the distribution of the number and sizes of bins created by different orderings proposed in the literature. The universal hitting sets ordering perform better than the other ordering in couple of ways. First, the number of bins created has a known bound, unlike other orderings that can create as many bins as there are *k*-mers (4^*k*^). In practice, this bound is also much lower than the actual number of bins created by other orderings. Second, the sizes of the bins are more uniform than with other orderings.

Based on this analysis of the performance of the minimizers algorithm, we advise bioinformatics tool authors to use the winnowing scheme with an appropriate universal hitting set in their application if possible, and a random ordering otherwise, in lieu of the default lexicographic ordering.

## 2 Approach

We will first give an overview of the results in this paper. Formal proofs for the results mentioned in this section are presented in subsequent sections.

In the original papers on minimizers and the winnowing schemes, the density is computed by considering any window of *w* + 1 consecutive *k*-mers. They make use of the following hypothesis:

**Hypothesis 1**. *Every k-mer in a* (*w + 1*)*-long window has an equal probability of 1/*(*w + 1*) *of being the smallest k-mer*.

Although not strictly true in practice, this hypothesis is reasonable and reflects reality accurately when using a randomized ordering. We define the *density factor d_f_* of a *k*-mer selection scheme *A* as the density times (*w* + 1), i.e.,

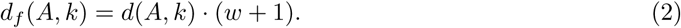

Assuming that Hypothesis 1 holds, we can rephrase the theorems of Schleimer *et al*. and Roberts *et al*. as

**Theorem 1** (Schleimer et al. [2003], Roberts et al. [2004b]). *Under Hypothesis 1, the density factor of the minimizers is d_f_* = 2.

A *universal k-mer hitting set* for given *k* and *w* is a set *U_k,w_* of *k*-mers such that every window of *w* consecutive *k*-mers must contain an element from *U_k,w_*. In Orenstein et al. [2016], we introduced universal hitting sets, and provided an efficient algorithm for constructing a compact universal set. These universal hitting sets do not need to contain all possible *k*-mers, and typically contain only about 1/*k* of all the *k*-mers. Here, we connect the ideas of universal *k*-mer hitting sets with minimizers by creating an ordering where the elements of *U_k,w_* are the smallest elements. Because every window contains an element of *U_k,w_* and these elements are smaller than other *k*-mers, only elements of *U_k,w_* can be selected as minimizers. Consequently, *k*-mers that are not in *U_k,w_* have probability 0 of being selected as minimizers and Hypothesis 1 does not hold anymore.

We now consider a refined hypothesis.

**Hypothesis 2**. *If a (w + 1)-long window contains j k-mers from a universal hitting set, each of these k-mers has an equal probability of 1/j of being the smallest k-mer.*

Motivated by the proof of Theorem 3 (see below), we define:

**Definition 2**. *The sparsity of a universal hitting set U_k,w_ **SP**(U_k,w_), is the proportion of (w+1)- long windows containing only one k-mer from U_k,w_.*

Assuming that Hypothesis 2 holds, we obtain a new estimation of the density factor.

**Theorem 3**. *Under Hypothesis 2, the density factor for the minimizers scheme with universal hitting sets U_k,w_ is*

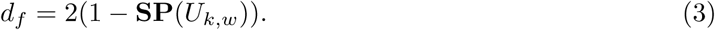

Note that when using the scheme we propose below, Hypothesis 2 provides no practical constraint on the use of minimizers in applications. Since any window will contain an element of *U_k,w_* by definition, any application assuming Hypothesis 1 can be made to work under Hypothesis 2 through the use of the appropriate *k*-mer ordering.

The expected density of the minimizers scheme for a given ordering is defined by an expectation over the set of all random sequences. So the density of minimizers computed for a given sequence might differ from the expected density. On the other hand, we will show how the expected density can be computed exactly by computing the density of minimizers on a particular sequence.

**Lemma 4**. *The expected density of a minimizers scheme with parameters k and w is equal to the density of minimizers on any de Bruijn sequence of order k + w*.

We also show that Theorem 3 provides a good approximation to the density for minimizers with universal hitting sets and exhibit counterexamples to the Schleimer *et al*. conjecture.

**Theorem 5**. *The density factor of a local scheme can be less than* 2.

The original intent of the minimizers is to select *k*-mers from a sequence as uniformly and sparsely as possible. Theorem 5 shows that schemes based on universal hitting set get closer to that goal. Using these schemes can greatly improve the performance of bioinformatics applications (See Section 4).

## 3 Proofs and Mathematical Details

### 3.1 Density with random ordering

First, we succinctly reproduce the proof from Schleimer et al. [2003] and Roberts et al. [2004b] of Theorem 1.

**Theorem 1** (Schleimer et al. [2003], Roberts et al. [2004b]). *Under Hypothesis 1, the density factor of the minimizers is ∑d_f_ = 2*.

*Proof*. Define a *charged* window *W*_*i*_ of *w* consecutive *k*-mers starting at position *i* in a sequence *S* as a window such that the smallest *k*-mer is different in *W_i_* than in *W*_*i*–1_. That is, as we sweep through the sequence from left to right, we charge the left-most window in which a given fingerprint is first selected. The number of fingerprints selected by the winnowing scheme is equal to the number of windows that are charged. Define the random variable *X_i_* to be 1 if *W_i_* is charged and 0 if not. Then the expected density can be written *d* = E[∑_*i*_*X*_*i*_]/*n* = ∑_*i*_ E[*X*_*i*_]/*n*, where *n* = |S| − *k* + 1 is the number of *k*-mers in the sequence.

Consider the larger window *W_i_^’^* of *w* + 1 *k*-mers starting at position *i* − 1 (*W*_*i*_ ⊂ *W_i_^’^*), and the smallest *k*-mer *m* in *W_i_^’^* (see Figure 1). If *m* starts at position *i* − 1, then *W*_*i*_ must be charged, as the previous fingerprint is now outside of the window *W*_*i*_. If *m* starts at position *i* + *w* – 1, then *W_i_* must be charged, as the new smallest *k*-mer just arrived in *W_i_*. In all other cases, *W*_*i*_ and *W*_*i*–1_ selected the same fingerprint and *W*_*i*_ is not charged. Assuming that the minimum *m* has equal probability to be in any of the positions [*i* − 1, *i* + *w* − 1], then E[X_*i*_] = P[*X_i_* = 1] = 2/(*w* + 1). So *d* = 2/(*w* + 1) and *d_f_* = 2.

**Figure 1:**
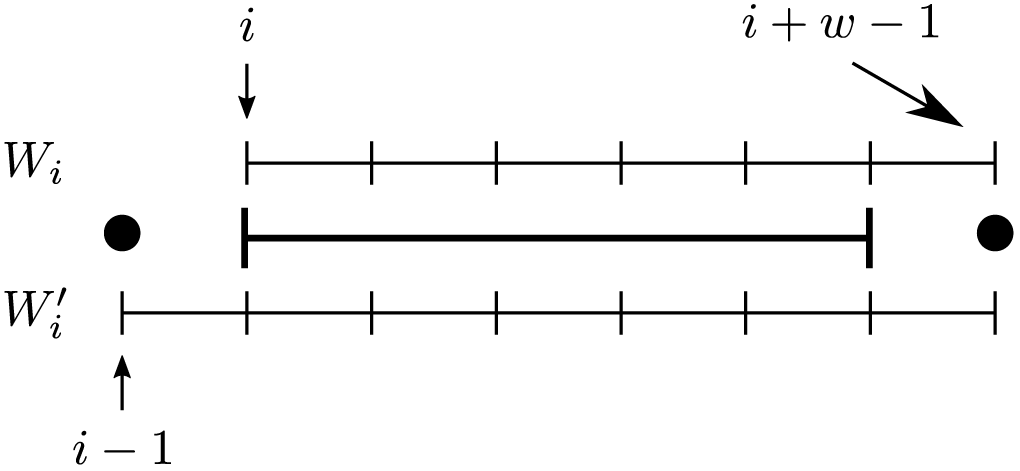
Windows *W*_*i*_, starting at position *i*, and window *W_i_^’^* starting at position *i* − 1. There are 3 different qualitative cases for the start position of the smallest *k*-mer *m: i−1* (left dot), *i*+*w*−1 (right dot) or in the range [*i,i+w−2*].

### 3.2 Computing the density

Even though the density is defined as an expectation over all possible random sequences where the bases are uniformly and independently chosen, for small values of *k* and *w*, it is possible in practice to compute the exact value of the expected density.

A *de Bruijn sequence* [de Bruijn, 1946] of order *k* on the alphabet Σ is a cyclic sequence that contains every possible *k*-mer as a substring exactly once and has length σ^*k*^, where σ is the number of symbols in the alphabet.

**Lemma 4**. *The expected density of a minimizers scheme with parameters *k* and *w* is equal to the density of minimizers on any de Bruijn sequence of order *k* + w*.

Proof. As seen in the proof of Section 3.1, whether a window *W_i_* is charged depends only on the sequence of the (*w* + 1)-long window *W_i_^’^*. Under the assumption that the input sequence is random, each (*w* + 1)-long window has a probability of 1/σ^*w+k*^. Hence, the density is the number of (*w* + 1)-long windows *W^’^* that cause the window *w* to be charged, divided by σ^*w+k*^. By computing the density on increasingly long random input sequences, the density would converge to the expected density as the proportion of the (*w* + 1)-windows converge to 1/σ^*w+k*^.

A de Bruijn sequence of order *k + w* contains every one of the σ^*w+k*^ possible (*w* + 1)-windows of *k*-mers exactly once (since it contains every (*w+k*)-long sequence exactly once). In other words, this de Bruijn sequence has all the (*w* + 1)-windows in exactly the desired proportion. Therefore, we can compute the expected density exactly by computing the density on the de Bruijn sequence of order *w + k*.

### 3.3 Universal hitting set ordering

The *de Bruijn graph G_k_* on a fixed alphabet Σ of size *σ* is a directed graph whose vertices are all the *k*-mers and a directed edge *(u, v)* represents a (*k*+1)-mer with *u* as its prefix and *v* as its suffix. A *universal hitting* set for the parameters *k* and *w* is a set of *k*-mers, *U_k,w_* so that any string of length *w + *k* −* 1 over Σ contains a substring from *U_k,w_*. Equivalently, every walk of length *w* over the nodes in the de Bruijn graph *G_k_* contains at least one *k*-mer from *U_k,w_*. In other words, it is a set of nodes that covers (has non-empty intersection with) all the length-*w* walks. (Here and throughout, the length of a walk is the number of vertices along it.) The set *V* (*G_k_*) of all the nodes of *G_k_* is trivially a universal hitting set, showing that the concept is well defined. In Orenstein et al. [2016], a heuristic is given to generate smaller universal hitting sets. The existence of such sets with certain desired properties is further discussed in Section 3.5.

Given a universal hitting set *U_k,w,_* we define an ordering <*_U_k,w__* on the *k*-mers as follows: u <*U_k,w,_ v* if *u* ∈ *_U_k,w__* and *v* ∉ *U_k,w_*; otherwise, *u <U_k,w_ v* if *u* is less than *v* in lexicographical order. In other words, the ordering makes the elements of the universal hitting set the smallest elements, and uses lexicographic order within the universal hitting set and its complement.

The most important property of this ordering is that, in each window of *w* consecutive *k*-mers, the smallest element must be an element of *U_k,w_*.

### 3.4 Density with universal hitting sets

With the ordering defined by a universal hitting set, we use the more appropriate Hypothesis 2 to prove the following theorem.

**Theorem 3.** *Under Hypothesis 2, the density factor for the minimizers scheme with universal hitting sets U_k,w_ is*

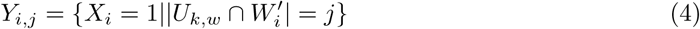

*Proof*. We adapt the proof of Section 3.1 to the ordering <*U_k,w_*. With this ordering, only the elements of *U_k,w_* are selected as fingerprints, and the number of such elements in the (*w* + 1)-long window varies between 1 and *w* + 1. We can compute the probability that a window is charged depending on how many elements from *U_k,w_* it contains. Let’s define the event *Y*_*i,j*_ that the window *W_i_* is charged, given that there are *j* elements of *U_k,w_* in the longer window *W_i_^’^* that contains it.

Then:

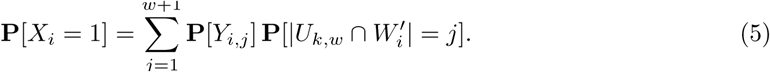

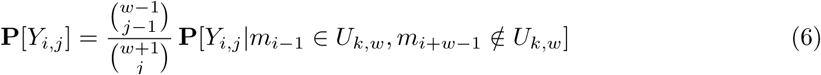

We can further condition the event *Y_i,j_* on whether the *k*-mers *m_i–1_* and *m_i+w–1_*, respectively at positions *i–1* and *i+w–1*, are in the universal hitting set. In the following, we use the convention that 
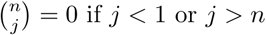
.

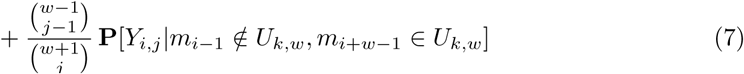

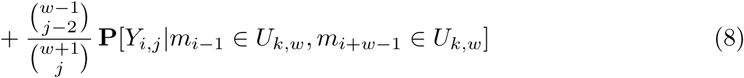

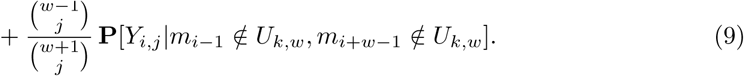

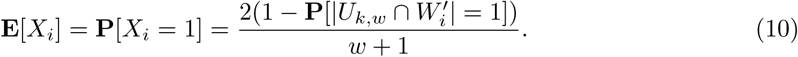

The factor 
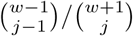
in (6) expresses the number of ways to place the *j* – 1 *k*-mers of *U_k,w_* in the *w* – 1 positions available (from *i* to *i* + *w*), given that *m_i–1_* is in *U_k,w_* and *m_i_+ *w* –1* is not in *U_k,w_*. The other factors are analogous.

We assume, based on Hypothesis 2, that all of the *j k*-mers from *U_k,w_*∩*W_i_^’^* have equal probability to be the smallest for the order <*U_k,w_*. Therefore the conditional probabilities equal 1/j for (6) and (7), 2/j for (8) and 0 for (9). Hence

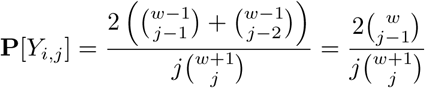

For 2 ≤ *j* ≤ *w* + 1, P[*Y*_*i,j*_] = 2/(*w* + 1). This is true in particular when *j = w* + 1, which corresponds to the case in the proof of Section 3.1. On the other hand, for *j* = 1, P[*Y*_*i*,1_] = 0. (This distinction motivates our definition of **SP**(*U_k,w_*).) This corresponds to the case where there is only one element from the universal hitting set in the (*w* + 1)-long window *W_i_^’^*. In that case, this element cannot be located at position *i* − 1 or *i + *w* –* 1 as that would leave a window of size *w* without any *k*-mer from *U_k,w_* which is impossible by construction. Hence, when *j* = 1, *W_i_* cannot be charged and **P**[Y_*i*,1_] has to be 0.

Finally, because 
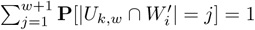
, we get from equation (5) that

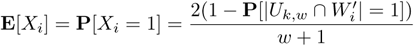

Under the assumption of random input sequence, the sparsity **SP**(*U_k,w_*) is precisely P[|*U_k,w_*∩*W_i_^’^*| = 1] and we have the main result *d_f_* = 2(1 − SP(*U_k,w_*)).

Equation (3) implies that if the universal hitting set *U_k,w_* is such that there exist (*w* + 1)-long walks in *G_k_* with only one element from *U_k,w,_* then the expected density factor is less than 2.

A universal hitting set *U_k,w_* that satisfies this condition, that is, **SP**(*U_k,w,_* > 0, will be called *w-sparse*. A *w*-sparse universal hitting set has density factor less than 2. See Section 3.7 for an example of *w*-sparse universal hitting set with low density.

Similarly to Section 3.2, the sparsity of a universal hitting set *U_k,w,_* can be computed exactly as the proportion of (*w* + 1)-long windows with only one *k*-mer from *U_k,w_* in the de Bruijn sequence of order *k + w*.

### 3.5 Existence of a sparse universal hitting set

We propose here a simple construction of *w*-sparse universal hitting sets to show their existence. This construction is not immediately useful in practice as the universal hitting sets it generates are large and have small sparsity (even though they are *w*-sparse). Different, more practical constructions are used when we discuss applications of this technique, in Sections 3.7 and 4.1.

Let *C* = (*c*_0_, …, *c*_ℓ−1_) be a simple cycle of length ℓ > 3 in *G*_*k*_, the de Bruijn graph of order *k* – 1. *C* is not necessarily an induced cycle, that is, it may have chords. Because *G_k_* is the line graph of *G*_*k* – 1_, the edges of cycle *C* induce a simple cycle *C_’_* of the same length in *G_k_*. That is, using indices modulo ℓ, node *c_i_^’^* of *G_k_* represents the edge (*c*_*i*_, *c*_*i*+1_) of *G_k_*–1 and *C*_*’*_ = (*c*_0_^*’*^,…,*c*^*’*^_ℓ−1_).

We claim that the cycle *C^’^* must be chordless. Suppose it is not the case, and there is an edge (*c^’^_i_*, *c^’^_j_*) where *j* ≠ *i* + 1. This implies that in *G_k_*–1 the head of edge *(c_i_, c_i+1_)* is equal to the tail of (*c*_*j*_, *c*_*j*+1_), that is *c*_*i*+1_ = *c*_*j*_. Because *j*≠*i*+1 and the cycle is simple, this is a contradiction.

Let *U*_*k*,*w*_ be all the *k*-mers, minus the nodes of *C′* unless their index is a multiple of *w*. That is *U*_*k,w*_ = (*V*(*G*_*k*_)\*C*^*’*^)⋃{*c*_*i*_^*’*^}_*w*|*i*_. Because *C′* is chordless, there is no cycle using only nodes not in *U_k,w_*. Also, by construction, there is no path of length *w* in *G_k_*\*U_k,w_*. Hence *U_k,w_* is a universal hitting set (though not a minimally sized one). By construction, for every *i* that is not a multiple of *w*, *W^’^_i_ = (*c*_*i*_^*’*^, *c*^*’*^_*i*+1_,…c^’^*_*i*+*w*_) contains only one element from *U_k,w,_* which implies that *U_k,w,_* is *w*-sparse.

So any cycle of length ℓ in *G_k−1_* induces a *w*-sparse hitting set for any *w*≤2(ℓ-1). In particular, any Hamiltonian cycle of *G_k−1_* induces a chordless cycle of length σ*^k−1^* in *G_k_* (these are the longest possible chordless cycles of *G_k_*). Therefore, for any given *k*, there are *w*-sparse universal hitting sets for all *w ≤ 2(σ^k−1^* − 1). Hence, there are sets for which **SP**(*U_k_w__*) is strictly greater than 0, indicating that sets with density factor < 2 exist.

Moreover, for small value of *w*, given that every cycle of length *w*/2 − 1 in *G_k−1_* induces a *w*-sparse universal hitting set in *G_k_*, there exists a large number of *w*-sparse universal hitting sets for the parameters *k* and *w*.

### 3.6 Universal hitting sets in random orderings

Consider again the winnowing scheme with random ordering, as we did in Section 3.1. In Section 3.1, it was assumed that all *w* +1 *k*-mers in the window *W_i_^’^* can potentially be selected as a fingerprint (Hypothesis 1). That assumption is likely not valid because of the existence and abundance of universal hitting sets. That is, even though the ordering of the *k*-mers is randomized, not all *k*-mers can be selected as minimizers with this ordering.

The following gives hints on why this is true. Consider a random ordering, or permutation, of the *k*-mers. Let *h(m)* be the index of the *k*-mer *m* in this permutation. In other words, *m* <_*h*_ *m^’^* ⇔ *h*(*m*) < *h*(*m^’^*). More precisely, consider a *k*-mer 
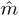
 that has a high index value in the permutation 
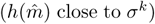
. If there exists a universal hitting set *U_k,w_* such that

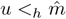
 for all *u ∈ U_k,w_* then the *k*-mer 
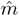
 cannot ever be selected by the winnowing scheme with that random ordering. This holds because every window of size *w* must contain a *k*-mer from *U_k,w_* which is less than 
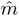
 in the ordering. Let *U* be the set of all universal hitting sets of minimum size and let *M* = min_U∈*U*_{max_*m*∈U_ *h*(*m*)}. Then any *k*-mer with an index greater than *M* can never be selected.

As a test, we ran the minimizers scheme (*k* = 10, *w* = 10, binary alphabet) for 1000 different random orderings on a de Bruijn sequence of order *w + k*. For each ordering *R_i_*, we obtain a set of minimizers *M_i_*, *1 ≤ i ≤ 1000*. Because we used a de Bruijn sequence of order *w + k*, the set *M_i_* contains a *k*-mer from every possible *w*-window. Hence *M_i_* is a, possibly trivial, universal hitting set for parameters *k* and *w*. Even though it is possible for *M_i_* to be the set of all *k*-mers, this was never the case in our 1000 random orderings, and the sets *M_i_* contain on average 51% of the *k*-mers (see Table 1).

**Table 1:** Statistics on the sparsity and density factor of the universal hitting sets generated by random ordering, the DOCKS universal hitting set and the lexicographic ordering. The computation is done with *k* = 10, *w* = 10 on a binary alphabet. The values for the random ordering are averages over 1000 different randomized orderings. *d_f_* is the density factor and *d_f,**SP**_*; is the density factor estimated given the sparsity of the set by equation 3. The difference between these numbers is due to the imperfect nature of Hypotheses 2.

| Ordering | $\mathbf{SP}(U_{k,w})$<br>% | $\vert U_{k,w}\vert /\sigma^k$<br>% | $d_f$ | $d_{f,\mathbf{SP}}$ |
| --- | --- | --- | --- | --- |
| random | 0.07 | 51 | 1.999 | 1.998 |
| DOCKS | 13.3 | 21 | 1.737 | 1.733 |
| lexicographic | 0.00 | 100 | 2.236 | 2.000 |

Furthermore, we computed the sparsity **SP**(*M_i_*) of the sets *M_i_*. In every case, the sparsity was small (average of 0:07%, see Table 1), but always greater than 0. Therefore, every observed universal hitting set *M_i_* is *w*-sparse. In other words, a random ordering implicitly defines a universal hitting set, and empirically this set is *w*-sparse in all randomized orderings we tested.

### 3.7 Density factor of various ordering schemes

Orenstein et al. [2016] introduces the concept of universal hitting sets and provides a heuristic algorithm, called *DOCKS* (Design Of Compact *K*-mer Sets), to generate small (although not necessarily optimal) universal hitting sets. Table 1 reports the actual density factor, computed as explained in section 3.2, of the winnowing scheme using three different orderings: randomized, DOCKS universal hitting sets and lexicographic. Because the densities are computed on a de Bruijn sequence of order *k* + *w* (binary alphabet, *k* = 10 and *w* = 10), the density computed is equal to the expected density.

In Schleimer et al. [2003], the authors prove a lower bound of 1.5, as well as a slight improvement of 1.5 + 1/2*w*, on the density factor for any local scheme. In addition, they conjecture that 2 is in fact the lowest possible density factor. The density factor of 1.737 obtained for DOCKS disproves this conjecture, but is not equal to the lower bound of 1.5 + 1/2*w* = 1.55, leaving a gap between the empirical results and the best known lower-bound.

The density factor estimation is obtained from the sparsity using equation 3. For the random and DOCKS orderings, this formula provides a good estimation of the density factor. The average sparsity of the random ordering is very low but not 0.

The DOCKS ordering, on the other hand, has a relatively high sparsity, which accounts for the low density factor. Also, the number of *k*-mers that are minimizers picked by the random and DOCKS orderings is significantly less than the total number of 10-mers.

The situation is very different for the lexicographic ordering, which selected all possible 10-mers and whose sparsity is 0. The estimation of its density factor from its sparsity is erroneous. This means that in addition to not being *w*-sparse, the lexicographic ordering does not satisfy Hypothesis 1. This is most likely due to the homo-polymer runs.

## 4 Application: binning

### 4.1 Binning DNA sequences

We consider the application of binning subsequences of a large DNA sequence, for example, as is done in *k*-mer counters such as KMC2 [Deorowicz *et al*., 2015] and MSPKmerCounter [Li and Yan, 2015]. These applications compute the number of occurrences of each *k*-mer in a DNA sequence. They are disk-based programs: first the input sequence is split into bins written to files on disk, then *k*-mer occurrences are computed in each bin independently. The number and size of bins considerably affects the running time of these programs.

For example, suppose we count L-mers in the human genome. Typically, for this application, 16 ≤ *L* ≤ 30. Following KMC2, we set *k* = 7 and *w* = *L* − *k* + 1, and we bin together every subsequence that shares a minimizer for parameters *k* and *w*. Then, it is possible to count the L-mers in each bin independently and in parallel to obtain the desired counts.

Ideally we would like a large number of bins, to allow for good parallelism, but not too large, to avoid extra overhead of creating and writing to many small files. Say at least a few hundred bins, but no more than a few thousand. For example, KMC2 uses a heuristic to merge in the same file the smaller bins so as to create at most 2000 files. Also, we would like the amount of data in each bin to be roughly the same. The lexicographic ordering for example generates a much larger bin corresponding to the homo-polymer of A. In particular, we would like the size of the largest bin to not be too large compared to the average bin size.

We next look at different orderings used in bioinformatics applications and their impact on the number of bins generated and the distribution of the data between the bins.

### 4.2 Summary of heuristic orderings used in practice

The following ordering was proposed by the authors of the minimizers technique in the implementation of the UMD Overlapper [Roberts *et al*., 2004a]. They “assign the values 0, 1, 2, 3 to C, A, T, G, respectively, for the odd numbered bases of *k*-mers and assign 0, 1, 2, 3 to G, T, A, C, respectively, for the even numbered bases.”

In the *k*-mer counter KMC2 [Deorowicz *et al*., 2015], the authors use the lexicographic order but ban a subset of *k*-mers from being minimizers. They use as minimizers *k*-mers that “do not start with *AAA*, neither start with *ACA*, neither contain *AA* anywhere except at their beginning”. This effectively creates a (non-optimally sized) universal hitting set (provided that the homopolymer *AA*…*A* is preserved).

In Kraken [Wood and Salzberg, 2014], the authors XOR the *k*-mer with a random value, before using lexicographic comparison. It is a form of randomization.

Minimap [Li, 2016] uses randomization as well, by employing a particular invertible hashing function.

### 4.3 Densities using various heuristics including a universal *k*-mer-based scheme

Figure 2 shows the distribution of the distance between consecutive minimizers, for *k* = 7 and *w* = 11, and Table 2 has statistics on these distributions. The distribution is computed for a de Bruijn sequence of order 18 = *k + w*, as explained in Section 3.2, and on the full sequence of the human genome (with all ambiguous bases and *N*s removed). Ideally, we would like the selection of *k*-mers to be as sparse as possible and, therefore, the distribution to be skewed toward larger distances.

**Figure 2:**
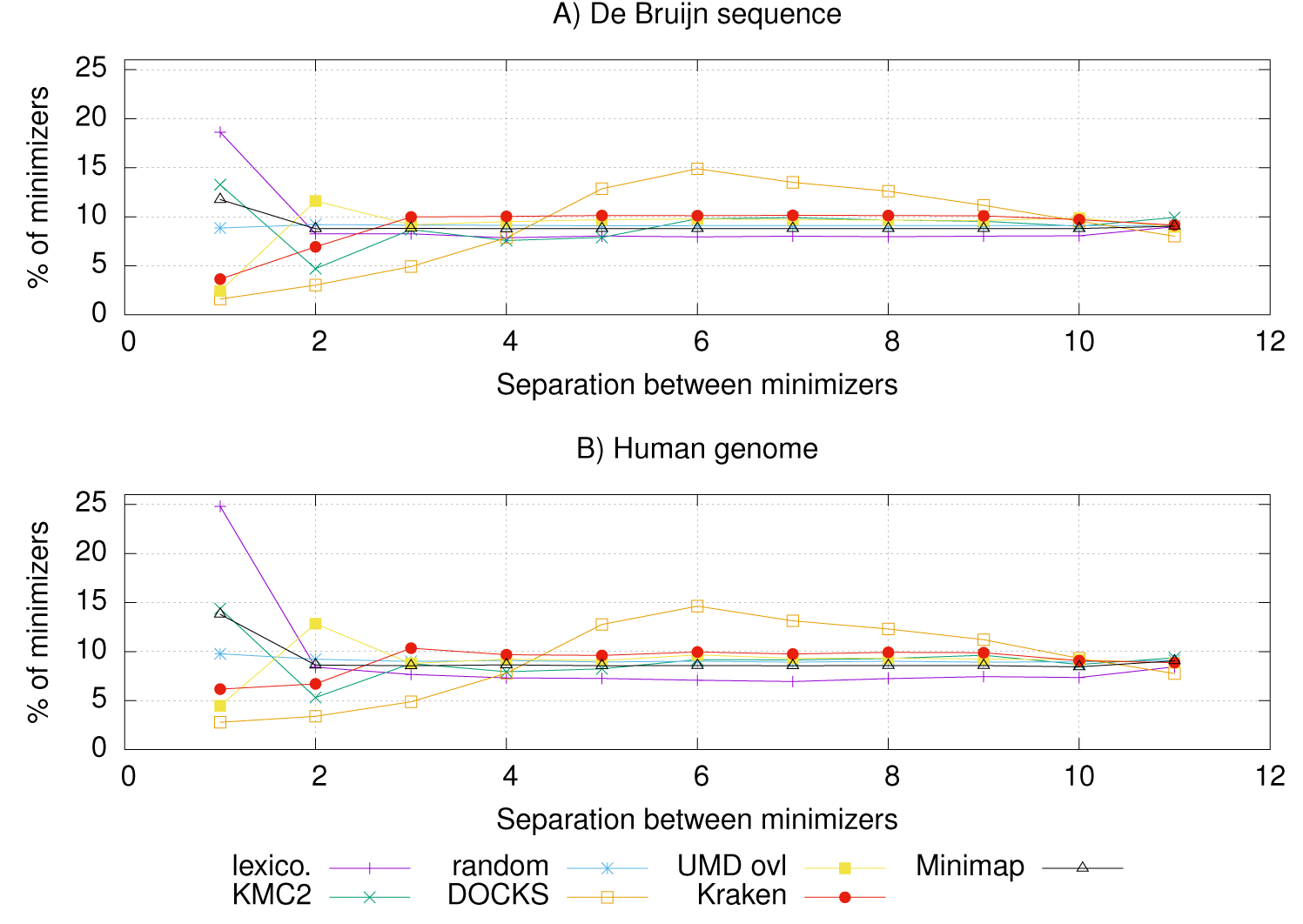
Distribution of the separation between minimizers for *k* = 7 and *w* = 11 on DNA sequences. (A) Results on a de Bruijn sequence of order *w* + *k*. (B) Results computed on the human reference genome (hg19). Each line represents a different minimizer scheme using a different ordering. Note that previous heuristic orderings all behave like the randomized orderings (uniform distribution) except for separation of 1 and 2. The universal *k*-mer ordering computed by DOCKS has a noticeably different distribution, with a mode and a higher mean.

**Table 2:** Statistics on the distribution of the distances between minimizers in Figure 2. The table reports the density factor (*d*_*f*_), the mean distance between minimizers (mean ± stdev) and the percentage of selected *k*-mers that are consecutive or separated by one base (low sep.). These were computed on a de Bruijn sequence (dbg) and on the human genome sequence (human).

| | Ordering | $d_f$ | distance<br>mean ± stdev | low sep.<br>% |
| --- | --- | --- | --- | --- |
| dbg | lexico. | 2.18 | 5.5 ± 3.4 | 27 |
| | random | 2.00 | 6.0 ± 3.2 | 18 |
| | Minimap | 2.05 | 5.9 ± 3.2 | 21 |
| | KMC2 | 1.97 | 6.1 ± 3.2 | 18 |
| | UMD Ovl | 1.91 | 6.3 ± 3.0 | 14 |
| | Kraken | 1.88 | 6.4 ± 2.9 | 11 |
| | DOCKS | 1.75 | 6.9 ± 2.5 | 4.6 |
| human | lexico. | 2.34 | 5.1 ± 3.4 | 33 |
| | random | 2.02 | 6.0 ± 3.2 | 19 |
| | Minimap | 2.09 | 5.8 ± 3.3 | 22 |
| | KMC2 | 2.02 | 5.9 ± 3.3 | 19 |
| | UMD Ovl | 1.97 | 6.1 ± 3.1 | 17 |
| | Kraken | 1.93 | 6.2 ± 3.0 | 13 |
| | DOCKS | 1.77 | 6.7 ± 2.6 | 6.2 |

The overall behavior for the different orderings does not change quantitatively between the de Bruijn sequence and the DNA sequence, showing that in the limit, the expected density computed on a de Bruijn sequence is a good proxy for performance on a biological sequence. The distribution of the randomized ordering is close to a uniform distribution. All the orderings succeeded in their stated goal of reducing the number of cases where minimizers have a low separation (e.g. *k*-mers that are consecutive or separated by one base) compared to the lexicographic ordering. For larger separation between minimizers, the distribution for all orderings except DOCKS are very similar to that of the randomized ordering.

The distribution for the DOCKS ordering is markedly different, with a mode at 6 (which is (*w* + 1)/2) and is generally skewed toward larger separation. The percentage of minimizers with low separation is the lowest among all orderings. This provides strong evidence that using a universal *k*-mer-based ordering can reduce the number of minimizers selected.

### 4.4 Distribution of bins created by heuristic and universal *k*-mer based schemes

Table 3 reports statistics on the number and sizes of the bins created using various orderings. The DOCKS based ordering creates far fewer bins than the other orderings, and the ratio between the largest bin and the average size is smaller. The DOCKS distribution of bin sizes is closer to the stated goals of not having too many bins, or any bins that are too large. These results indicate that if these tools adopted a universal *k*-mer ordering, their runtime and memory usage performance would be significantly improved.

**Table 3:** Statistics on the distribution of the bin sizes. The table reports the number of bins created (# bins), the average bin size (avg size, in million of bases) and the ratio of the largest bin size to the average (max ratio). These were computed on a de Bruijn sequence (dbg) and the human genome sequence (human).

|  | Ordering | # bins | avg size<br>(million bases) | max ratio |
| --- | --- | --- | --- | --- |
| dbg | lexico. | 16 384 | 4.19 | 367 |
|  | random | 12 003 | 5.73 | 9.32 |
|  | Minimap | 13 267 | 5.18 | 10.4 |
|  | KMC2 | 12 370 | 5.56 | 166 |
|  | UMD Ovl | 13 108 | 5.24 | 274 |
|  | Kraken | 12 502 | 5.5 | 210 |
|  | DOCKS | 4063 | 16.9 | 7.44 |
| human | lexico. | 16 285 | 0.175 | 96.4 |
|  | random | 11 388 | 0.251 | 38.3 |
|  | Minimap | 12 280 | 0.233 | 69.5 |
|  | KMC2 | 12 287 | 0.233 | 74.5 |
|  | UMD Ovl | 11 015 | 0.259 | 26.1 |
|  | Kraken | 10 389 | 0.275 | 26.3 |
|  | DOCKS | 4046 | 0.706 | 23.5 |

## 5 Discussion

We introduced a new analysis of the performance of the winnowing scheme based on the sparsity of an ordering. In addition, we showed how to construct orderings from small universal hitting sets that outperform randomized and previously designed orderings. While these new orderings perform well on uniformly distributed sequences and actual DNA sequences, many questions of theoretical and practical interest remain.

First, several orderings based on universal hitting sets beat the conjectured lower bound on the density factor of 2, but did not achieve 1.5 + 1/2*w*. The question of the actual lower bound is therefore open. Second, every winnowing local scheme considered here is based on an ordering of the *k*-mers. But there exists local schemes that are not based on an ordering. Can the minimum density always be achieved with an ordering? Third, although the performance was similar between the uniformly distributed input sequence and DNA sequence, can we improve the winnowing scheme by taking into account the actual distribution of the bases in the input?

Fourth, the construction of small universal hitting sets with DOCKS is based on a heuristic and is not guaranteed to be optimal. As it stands, it is also intractable for large *k*. Can we design an efficient algorithm to construct some (or all) of the optimal universal hitting sets? Or, can we design optimal orderings, even though the actual corresponding universal hitting set is never explicitly constructed?

All of the work in this paper considers a de Bruijn graph. In many situations, it makes sense to consider a *k*-mer and its reverse complement to be equal. The above schemes can be used by replacing every *k*-mer with its canonical representation (i.e., the smallest of a *k*-mer and its reverse complement). But this is likely not optimal. Define the RC *de Bruijn graph* from a de Bruijn graph where nodes representing reverse complements are identified and parallel edges are collapsed into a single edge. A fifth question arises in this setting: can an equivalent to universal hitting sets, and orders, be defined so as to create better winnowing schemes when reversed complemented *k*-mers are identified? How much can we transfer of what we already know on de Bruijn graphs to RC de Bruijn graphs?

## 6 Conclusion

We provided a novel theoretical framework to estimate the density of minimizer procedures. This framework explains why it is possible for *w*-sparse local winnowing schemes to have lower density factor than 2. In addition, we provided a practical method to create from small universal hitting sets, such as those generated by DOCKS, *w*-sparse winnowing schemes that achieve density factors below 2. These two results combined answer negatively a standing conjecture of Schleimer *et al*.

We showed that the lexicographic ordering has the worst behavior while the different orderings used in current software tools perform similarly to randomized orderings, which is slightly better. We showed that universal hitting set based schemes can perform substantially better than random orderings in practice. Therefore, for the development of bioinformatics software using minimizers, we suggest using an ordering based on small universal hitting sets such as those created by the DOCKS algorithm, or, if not practical, randomized orderings. We showed that for binning applications, doing so would lead to a significant improvement in computational efficiency.

## Acknowledgement

This research is funded in part by the Gordon and Betty Moore Foundation’s Data-Driven Discovery Initiative through Grant GBMF4554 to C. K., by the US National Science Foundation (CCF-1256087, CCF-1319998) and by the US National Institutes of Health (R01HG007104). D. P. was supported in part by a Ph. D. fellowship from the Edmond J. Safra Center for Bioinformatics at Tel-Aviv University. R. S. was supported in part by the Israel Science Foundation as part of the ISF-NSFC joint program 2015-2018.

